# dGPredictor: Automated fragmentation method for metabolic reaction free energy prediction and *de novo* pathway design

**DOI:** 10.1101/2021.03.15.434460

**Authors:** Lin Wang, Vikas Upadhyay, Costas D. Maranas

## Abstract

Group contribution (GC) methods are conventionally used in thermodynamics analysis of metabolic pathways to estimate the standard Gibbs free energy change (*Δ*_*r*_*G*′^*o*^) of enzymatic reactions from limited experimental measurements. However, these methods are limited by their dependence on manually curated groups and inability to capture stereochemical information, leading to low reaction coverage. Herein, we introduce an automated molecular fingerprint-based thermodynamic analysis tool called dGPredictor that enables the consideration of stereochemistry within metabolite structures and thus increases reaction coverage. dGPredictor has a higher prediction accuracy as compared to existing GC methods and can capture free energy changes for isomerase and transferase reactions, which exhibit no overall group changes. We also demonstrate dGPredictor’s ability to predict the Gibbs free energy change for novel reactions and seamless integration within *de novo* metabolic pathway design tools such as novoStoic. This enables performing a thermodynamic analysis for synthetic pathways, thus safeguarding against the inclusion of reaction steps with infeasible directionalities. To facilitate easy access to dGPredictor, we developed a graphical user interface to predict the standard Gibbs free energy change for reactions at various pH and ionic strengths. The tool allows customized user input of known metabolites as KEGG IDs and novel metabolites as InChI strings (https://github.com/maranasgroup/dGPredictor).

**Author summary:** The genome-scale metabolic networks consist of a large number of biochemical reactions interconnected in a complex system. The standard Gibbs free energy change is commonly used to check for the feasibility of enzyme-catalyzed reactions as thermodynamics plays a crucial role in pathway design for biochemical synthesis. The group contribution methods using expert-defined functional groups have been extensively used for estimating standard Gibbs free energy change with limited experimental measurements. However, current methods using functional groups have major issues that limit its ability to cover all the metabolites and reactions as well as the inability to consider stereochemistry leads to erroneous estimation of free energy that undergoes only stereochemical change such as isomerases. Here, we introduce a molecular fingerprint-based thermodynamic tool dGPredictor that enables stereochemistry in metabolites and thus improves the reaction coverage with higher prediction accuracy compared to current GC methods. It also allows the ability to predict free energy change for novel reactions which can aid the *de novo* metabolic pathway design tool to ensure the reaction feasibility. We apply and test our method on reactions in the KEGG database and isobutanol synthesis pathway. In addition, we provide an open-source user-friendly web interface to facilitate easy access for standard Gibbs free energy change of reactions at different physiological states.

## Introduction

Thermodynamics determines both the directionality and efficiency of enzymatic reactions, thereby dictating metabolic phenotypes for product synthesis in microbial engineering platforms. For example, the beta-oxidation pathway can be reversed because the overall standard Gibbs free energy change in the reverse direction becomes negative when utilizing ferredoxin as the reducing equivalent [1]. This engineered pathway enables the production of higher-chain linear alcohols and fatty acids with greater efficiency. The biological methanogenic and acetogenic reduction pathways are highly efficient in converting CO_2_ to CH_4_ due to lower thermodynamic barriers compared to the corresponding geochemical pathways [2]. Thus, thermodynamic analysis is an important tool for assessing and selecting feasible heterologous metabolic pathways [3] and quantifying the thermodynamic driving force for biosynthesis in different production hosts using intracellular metabolomic data [4]. However, direct experimental measurements of standard Gibbs free energy change of reactions (*Δ*_*r*_*G*′^*o*^) are still limited to ∼600 enzymatic reactions cataloged in the Thermodynamics of Enzyme-catalyzed Reactions Database (TECRDB) [3]. Emerging isotopic labeling experiments such as deuterium-labeled studies [5] can directly quantify *Δ*_*r*_*G*′^*o*^ of enzymatic reactions but have so far been limited to central carbon metabolism thus necessitating the use of predictive computational frameworks.

In order to expand the prediction of *Δ*_*r*_*G*′^*o*^ for reactions lacking experimental measurements, group contribution (GC) methods were developed. They assume that the *Δ*_*r*_*G*′^*o*^ can be expressed as the sum of contributions from all functional groups (*Δ*_*g*_*G*^*o*^) (based on a predefined list of substructures [6]), which in turn were fitted from experimental data. Multiple linear regression is typically applied to determine each group’s contribution in a reaction by minimizing the mean squared error between the observed and estimated *Δ*_*r*_*G*^*o*^. Various group contribution methods were developed to improve the prediction accuracy [7]. Table 1 contains a brief description of existing methods for *Δ*_*r*_*G*′^*o*^ estimation.

**Table 1.** Existing methods for the prediction of standard Gibbs free energy of biochemical reactions.

| Algorithm/Method | Citation | Major contributions |
| --- | --- | --- |
| Group contribution | Mavrovouniotis et al. (1990) | Estimate free energy of formation by decomposing compounds into functional groups based on biochemical knowledge |
| Group contribution | Jankowski et al. (2008) | Free energy estimation for biochemical reactions in *Escherichia coli* metabolic network and improved previous method by introducing additional groups and interaction factors |
| Group contribution | Noor et al. (2012) | Improved last method by integrating known thermodynamic data for molecules when available<br>Considered pseudoisomers of molecules at different protonation levels to accounts for the effect of pH and ionic strength |
| eQuilibrator | Flamholz et al. (2012) | User-friendly web-based platform based on previous group contribution method by Noor et al. |
| Component contribution | Noor et al. (2013) | Prioritize on reactant contribution over group contribution, which directly uses the formation energy when available |
| Updated group-contribution | Du et al. (2018) | Improved previous methods using entropy and enthalpy information<br>Considered influence of magnesium binding with metabolites and temperature |

Despite the extensive efforts to improve group contribution predictions, the approach itself suffers from many inherent limitations [7]. The expert-defined groups provide incomplete coverage leading to (i) metabolites that cannot be decomposed, and thus their *Δ*_*f*_*G*^*o*^ cannot be estimated, (ii) an assignment of zero for *Δ*_*r*_*G*^*o*^ of reactions with no group changes despite experimental values being non-zero (such as isomerase reactions), and (iii) large uncertainties in *Δ*_*r*_*G*^*o*^ for reactions that involve groups occurring sparingly in the training dataset.

Moving beyond simple molecular groups, more than 11,145 descriptors have been developed for extracting chemical information for describing Quantitative Structure-Activity Relationships (QSAR) [13]. Molecular descriptors such as molecular [14] and circular fingerprints [15] use fragments/moieties to represent substructure information as vectors, matrices, or other mathematical representations such as encoding group information in GC methods. They were applied extensively, alongside statistical and machine learning algorithms, to predict physical and (bio-)chemical properties (such as dissociation constant, viscosity, and toxicity) for drug design [16]. However, their application for *Δ*_*r*_*G*′^*o*^ estimation of enzymatic reactions has been limited to only a few studies [17,18]. The computation tool IGERS [17] predicts *Δ*_*r*_*G*′^*o*^ of a new reaction based on its similarity (calculated using 2D molecular descriptors) to reference reactions with experimental measurements. More recently, a machine learning algorithm using chemical fingerprints as a feature [18] showed improved prediction accuracy.

Chemical substructures have been used extensively to encode molecular information in many computational *de novo* pathway design tools [19]. Substructure changes in enzymatic reactions can be generalized as rules, thereby codifying *de novo* reactions to fill in missing chemical conversion steps. However, current *de novo* pathway design tools allow only *a posteriori* evaluation of thermodynamic feasibility of a proposed metabolic pathway as novel reaction steps are generally treated as being reversible [20]. Therefore, significant computational resources may be expended in generating pathways with one or more steps operating in a thermodynamically unfavorable direction. Even though tilting the relative reactant/product concentrations can maintain feasibility of the designed pathway, the imposed concentration ranges may not be physiologically viable. Furthermore, operating near thermodynamic equilibrium dramatically increases the required enzyme level, incurring a significant metabolic burden [10,21]. Thus, an automated approach for estimating the *Δ*_*r*_*G*^′*o*^ of all novel steps is required, which can also be easily embedded within pathway design tools such as novoStoic [22]. This would help refine *de novo* predictions by constraining the reaction steps/rules in only the thermodynamically feasible direction.

Herein, we developed a moiety-based automated fragmentation tool called dGPredictor for *Δ*_*r*_*G*^′*o*^ estimation of enzymatic reactions. Moieties are descriptors of the bonding environment of all non-hydrogen atoms in a chemical structure. They differ from functional groups, which are a set of connected atoms responsible for a shared characteristic such as chemical nomenclature, reactivity, and bond types (single, double, etc.) [23]. Multiple models were tested within dGPredictor using moiety descriptions of different spans and using both explainable linear regression and neural network-based nonlinear formalisms. Resultant prediction accuracies and overfitting potential were calculated as the mean squared error (MSE) over training data and median of mean absolute error (MAE) of leave-one-out-cross-validation (LOOCV) results, respectively, which were used as two metrics to compare the prediction accuracies for Δ _*r*_ *G*′^*o*^ estimates and facilitate the direct comparison with widely used GC method (specifically, component contribution (CC) [10]). dGPredictor improved the accuracy of prediction over the widely used CC method [10] by 78.76% (i.e., MSE over training data from TECRDB database (see Table 2)). It also led to an increase in the coverage of Δ_*f*_*G*′^*o*^ and Δ_*r*_*G*′^*o*^ estimation for metabolites and reactions present in the KEGG database by 17.23% and 102%, respectively, over GC by allowing for stereochemical considerations not captured in previous expert-defined chemical groups. Examining the sensitivity of model predictions to moiety definition revealed that moieties spanning a bonding distance of two are more prone to overfitting as compared to a one due to combinatorial explosion of unique moiety types (e.g., cross-validation MAE of 5.83 vs. 15.46 for bonding distance one and two, respectively). Moieties spanning distances one and two in a combined linear model were found to be less prone to overfitting than the corresponding non-linear variants (cross-validation MAE 5.48 vs. 7.27 for the best performing non-linear NN model). This is likely due to the relatively small size (i.e., 4,001 reactions and 673 metabolites) of the training dataset that does not seem to benefit from using a more extensive set of descriptors embedded in a non-linear modeling framework.

**Table 2.** Details of prediction accuracy and cross-validation scores for different regression models.

| Model | Mean squared error | cross-validation: Mean absolute error |
| --- | --- | --- |
| M1-linear | 38.30 | 5.83 |
| M2-linear | 24.60 | 15.46 |
| **M1,2-linear** | **9.60** | **5.48** |
| M1-nonlinear | 20.81 | 6.64 |
| M2-nonlinear | 6.92 | 14.69 |
| M1,2-nonlinear | 6.95 | 7.27 |
| **GC [10]** | **45.20** | **5.32** |

Next, we employed dGPredictor to aid metabolic pathway design by improving upon the current practice of considering *de novo* reactions as being always reversible, thereby necessitating additional scrutiny to ensure thermodynamic feasibility. We show that dGPredictor can estimate the Δ_*r*_*G*′^*o*^ of *de novo* reactions (i.e., involving novel metabolites) by utilizing their chemical structure information in the IUPAC International Chemical Identifier (InChI) strings [24]. Furthermore, pathway design tools can use the moiety changes in dGPredictor as reaction rules to ensure the thermodynamic feasibility.

## Results

### Automated fragmentation of metabolites

In contrast to the widely used GC methods that rely on expert-defined groups, we apply an automated fragmentation approach similar to Carbonell et al. [14] to encode novel molecules. dGPredictor classifies every (non-hydrogen) atom in a structure by assessing its bonding environment at a distance of one (M1 moieties) or two bonds (M2 moieties). We do not consider bonding distance zero (M0 moieties) and more than two (M2+ moieties), as M0 only encode the element associated with each atom and to avoid overfitting with M2+ moieties as the available regression dataset was relatively limited in size. We use (+)-epi-isozizaene as an example to demonstrate the chemical moieties-based metabolite descriptors used in dGPredictor. Fig 1A shows the seven chemical moieties that are generated from (+)-epi-isozizaene when stereochemistry is considered. The occurrence of the seven moieties is counted and used as the group/moiety incidence matrix *G*_*i,g*_ (shown in blue, where rows denote moieties, and the columns count the occurrence of each moiety in the chemical structure and corresponding Simplified molecular-input line-entry system (SMILES) representation). The moiety shown in orange was obtained from the SMILES annotation C[C@@H](C)C, where the chiral specification (i.e., “C@@H”) indicates the carbon atom is the tetrahedral center. The two different symbols, “@” and “@@” indicate anticlockwise and clockwise configurations, respectively. Without stereochemistry, (+)-epi-isozizaene can be decomposed into only five moieties (see Fig 1B). The moiety shown in green is now represented by the SMILES string CC(C)C without chiral specifications. Thus, including stereochemistry-based decomposition within dGPredictor allows incorporating moiety changes for reactants and products that differ only in stereochemistry (e.g., isomerases). Fig 1A and B highlight the possible loss of stereochemical information that occurs when not considering the two possible configurations around an asymmetric tetrahedral atom, which results in the same moiety description for both configurations.

**Fig 1.**
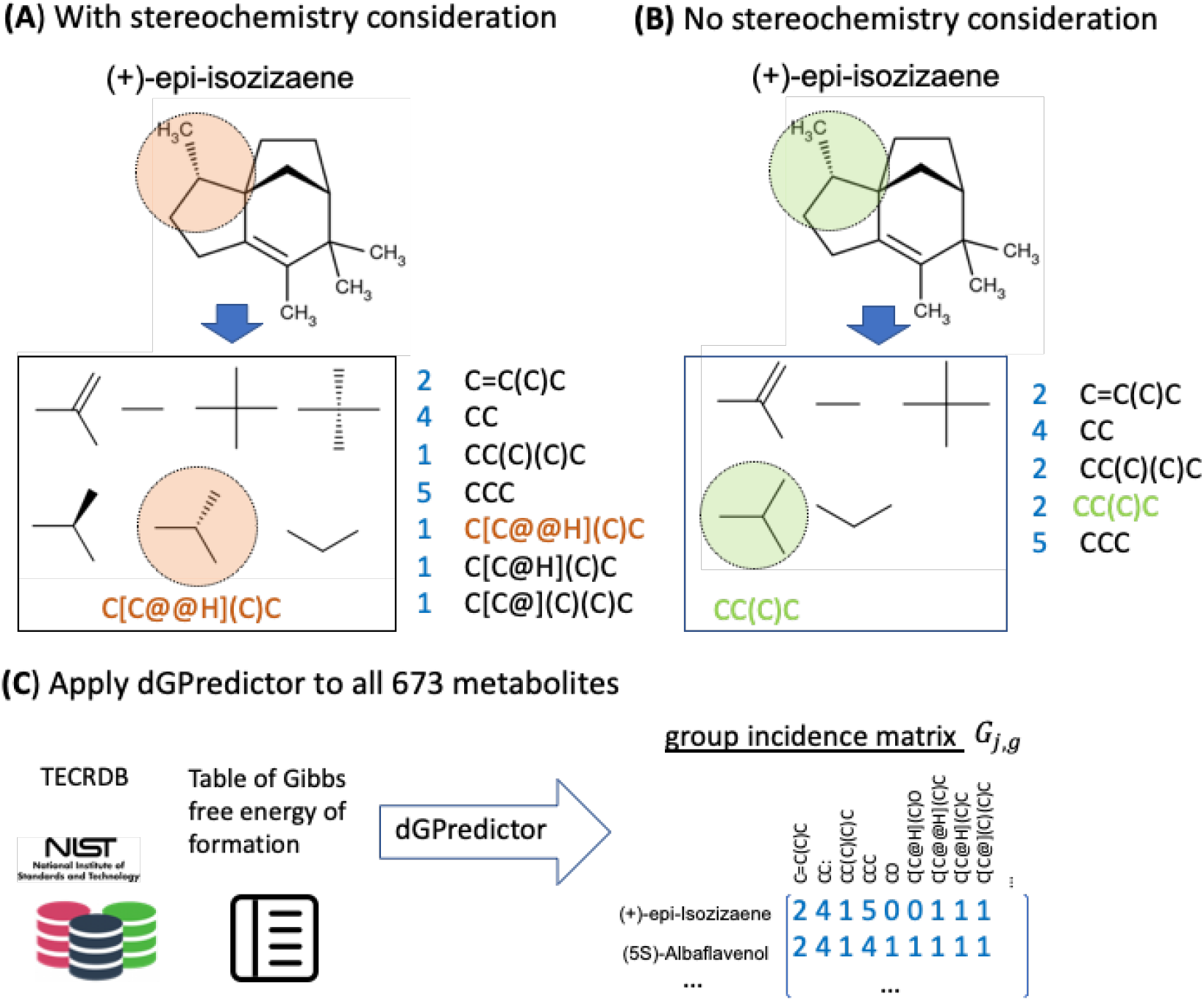
The substructures/chemical moieties generated by dGPredictor. The decomposition of (+)-epi-isozizaene (A) with stereochemistry consideration and (B) without stereochemistry consideration, and (C) group/moiety incidence matrix created by dGPredictor for all metabolites in TECRDB.

It is worth noting that the recent GC method, component contribution [10] cannot decompose (+)-epi-isozizaene due to the incomplete coverage of all substructures in CC. As shown in Fig 1, dGPredictor can encode metabolites with both very complex structures as well as very small chemicals such as hydroxylamine (NH_2_OH). We applied dGPredictor to construct the group/moiety incidence matrix for the 673 metabolites with experimental measurements in TECRDB (see Fig 1C). For moieties spanning a bonding distance of one and two, we generated 263 (S1 table 1) and 1,380 (S2 Table) members, respectively, using the RDkit python package (See Methods section ‘Automated fragmentation of metabolites’).

### Improved goodness of fit of Δ_*r*_*G*′^*o*^ by dGPredictor using an automated moiety classification method

We apply Bayesian ridge regression [25,26] and feedforward neural networks [27,28] to estimate the standard Gibbs free energy contributions of the moieties. Moiety decomposition and the regression models employed are described in detail in the Method section. Note that as experimental data was collected at different pH and ionic strength, a mixture of pseudoisomers (i.e., multiple structures with different protonation states) exists for each metabolite. We used the Inverse Legendre Transform (see details in Methods section ‘Pseudoisomer’) to standardize the experimental values of Δ_*r*_*G*^*o*^ at pH 7 and ionic strength 0.25 M [29]. Inverse Legendre Transform reduces the ensemble of pseudoisomers to a single pseudoisomer (i.e., the most abundant pseudoisomer at pH 7) and was used as the reference for regression analysis in this section. The difference between Δ_*r*_*G*^*o*^ calculated for a single pseudoisomer and the transformed Gibbs energy Δ_*r*_*G*′^*o*^ (i.e., Gibb’s energy of the ensemble) at a specific cellular condition is corrected as a function of the dissociation constant *pKa* for each pseudoisomer, pH, and the ionic strength when making predictions (see details in Methods section ‘Pseudoisomers’).

First, we applied Bayesian ridge regression to estimate the individual contribution of each moiety from available experimental measurements of Δ_*r*_*G*^*o*^. We chose to assess moieties associated with a bonding distance one and two to select the model with better prediction accuracy. In the remainder of the paper, we refer to the models of distance one and two as M1-linear (263 unique moieties for 4,001 reactions) and M2-linear (1,380 moieties for 4,001 reactions), respectively. In addition, we examined the efficacy of using moieties of both distances simultaneously within a single model referred to as the M1,2-linear (1,643 moieties for 4,001 reactions). Model prediction accuracy is defined as the mean squared error (MSE) over the entire training dataset. For the M2-linear model, MSE improved by 35.77% over the M1-linear (see Table 2), which is expected as the number of moieties increases >5-fold from 263 to 1,380. We found that combining moieties of radius one and two in model M1,2-linear further lowered the MSE by 60.97% compared to the M2-linear. Model overfitting potential in the cross-validation analysis is defined as the mean absolute error (MAE) 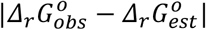 between the experimentally observed Gibbs free energy change of a reaction (i.e., 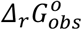 in the validation dataset) and the predicted value from the linear regression model trained without the experimental data of that particular reaction (i.e., 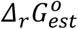 in the training dataset). This metric can be used to flag model overfitting by comparing MAE values between models – a larger MAE value on the validation data alongside a lower MSE value on the training data indicates overfitting. However, a leave-one-out cross-validation (LOOCV) analysis of all models revealed over-fitting by the M2-linear as compared to the other two linear models. We found a significant increase in MAE for M2-linear compared to M1-linear and M1,2-linear models, with a median MAE of 15.46, 5.83, and 5.48, respectively (see Table 2). The M1,2-linear model provides the lowest MSE on the training dataset with a value of 9.60 compared to 38.30 for M1-linear and 24.60 for M2-linear and a significantly lower overfitting potential. Therefore, we choose the M1,2-linear as the best among the three linear models. Do note that this should not be interpreted as a result that would hold universally. It is likely due to the small training set size (i.e., 4,001 reactions) compared to the total number of possible moieties.

### Introducing neural network-based nonlinear moiety contribution leads to a marginal increase in prediction performance

We next explored whether the inherent linearity of moiety contributions in the developed models limits their predictive capability. We used a feed-forward multi-layer neural network to allow for a nonlinear description of moiety contribution for predicting Δ_*r*_*G*^*o*^. Similar to linear models, three models with varying moiety spanning distance (same as linear regression models in the above section) were generated to determine the effect of moiety description on prediction accuracy - M1-nonlinear, M2-nonlinear, and M1,2-nonlinear. Similar to the linear models, Δ_*r*_*G*^*o*^ prediction accuracy (i.e., a lower MSE value) improved with increasing bonding distance (see Table 2). Models M2-nonlinear and M1,2-nonlinear were nearly identical in their prediction accuracy within the training datasets (MSE 6.92 vs. 6.95) and outperformed the M1-linear model (>3-fold higher MSE 20.81). The LOOCV MAE was lower for the M1,2-nonlinear model than the M2-nonlinear (7.27 vs. 14.69, see Table 2), indicating that the former is the best-performing nonlinear model.

A comparison among the six models (three linear and three nonlinear) suggests MSE values for all three nonlinear models were somewhat lower than their linear counterparts. However, a correspondingly higher LOOCV MAE revealed that the increased accuracy was likely due to overfitting. M1,2-nonlinear has a slightly lower MSE compared to the M1,2-linear (6.95 vs. 9.60). However, cross-validation scores showed that the M1,2-linear is significantly less susceptible to overfitting with MAE 5.48 vs. 7.27. This implies that the extra complexity and lack of interpretability associated with the M1,2-nonlinear model do not come with any appreciable increase in prediction performance. Since the MSE of both models is comparable, we chose the model that is least prone to overfitting as our overall best model, i.e., the M1,2-linear, and all subsequent results were computed using this model.

Finally, we assessed model performance between the automated decomposition scheme using moieties proposed herein and expert-based groups. We applied the same set of thermodynamics data compiled by Noor et al. [10] (available at https://github.com/eladnoor/component-contribution/) on dGPredictor and the GC method to enable direct comparison. dGPredictor using the M1,2-linear model outperformed GC, with a lower MSE (9.60 vs. 45.20) and a higher R^2^ score (0.9998 vs. 0.9989) (Fig 2A) estimated over the entire training dataset. This can be attributed to the underlying energy contribution of groups/moieties estimated by the two competing methods. As evaluated by Du et al. [7], the linear regression-based GC model often predicts unrealistically large |Δ_*g*_*G*^*o*^| for functional groups with limited representation in the training data which was observed in GC that predicted a |Δ_*g*_*G*^*o*^| value greater than 1,500 kJ/mol for two groups (Fig 2B), while the largest value in dGPredictor is ∼200 kJ/mol. dGPredictor mitigates this issue by applying regularization in the regression analysis. It must be noted that the median MAE from GC and dGPredictor is 5.32 kJ/mol and 5.48 kJ/mol respectively (Fig 2C) indicating similar overfitting potential.

**Fig 2.**
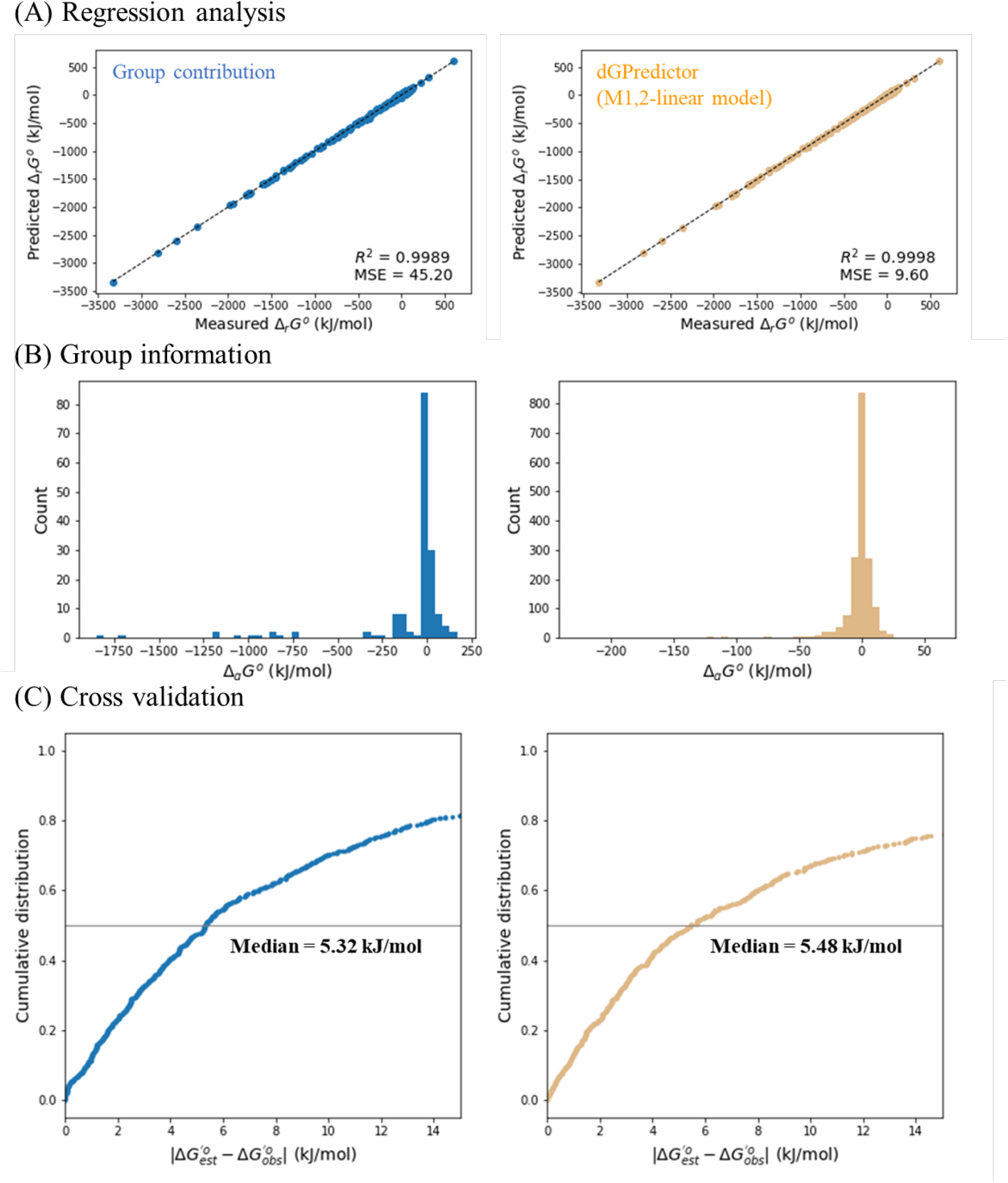
Comparison between group contribution method by Noor et al. [10] and dGPredictor (best model: M1,2-linear). (A) Comparison of regression analysis for the best dGPredictor model (M1,2-linear model) and previous group contribution method. It shows the improvement using two metrics mean squared error (MSE improved from 45.20 to 9.60) and R-squared (increased from 0.9989 to 0.9998). (B) Histogram denoting the estimated contribution of groups. dGpredictor group contribution estimates do not predict unrealistically high values, unlike group contribution. (C) leave-one-out cross-validation results, which indicates very close performance with the previous method with an improved MSE value.

### Increased coverage of metabolites and reactions by dGpredictor

A rigorous test of any predictive model lies in its ability to generalize over unseen data. Thus, we used the KEGG database [31] to test dGPredictor’s scope and prediction capability for reactions that are not included in the training dataset. This would also help evaluate the model’s ability to provide genome-scale coverage and estimate Δ_*r*_*G*^*o*^ for novel reactions when designing *de novo* biochemical pathways. The KEGG database consists of 15,278 metabolites and 7,053 reactions with chemical structure information defined by InChI strings. We first compared the ability of the GC method and the automated fragmentation method of dGPredictor to describe all metabolites present in the database as chemical groups/moieties. The GC method could describe 85.3% of the 15,278 metabolites, while dGPredictor succeeded in covering all metabolites (see Table 3). The GC method missed 14.7% metabolites due to them containing unique substructures which are not included in its expert-defined 163 groups. Fig 3A illustrates a few of these unique moieties and the corresponding dGPredictor decomposition. Most of the substructures that were not covered in GC methods decomposition involved bonds with N, P, and S atoms. Next, the *Δ*_*r*_*G*^*o*^ of reactions in the KEGG database was calculated using dGPredictor and GC. Metabolites from the TECRDB database [32] were decomposed using expert-defined GC groups and the automated moiety-based framework in dGPredictor and used to calculate Δ_*r*_*G*^*o*^. GC could successfully describe 33.8% of the database reactions, while dGPredictor could parameterize twice that number (i.e., 69.3%), indicating that the developed formalism can successfully generalize to provide larger reaction coverage.

**Table 3.**
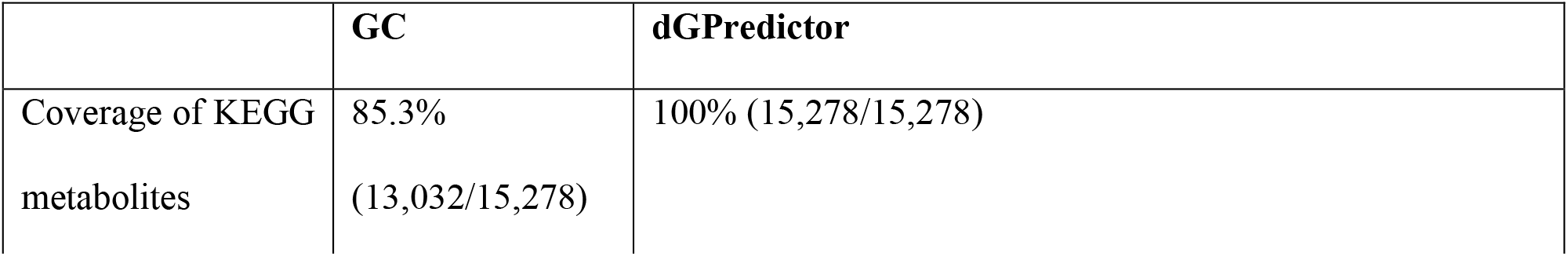
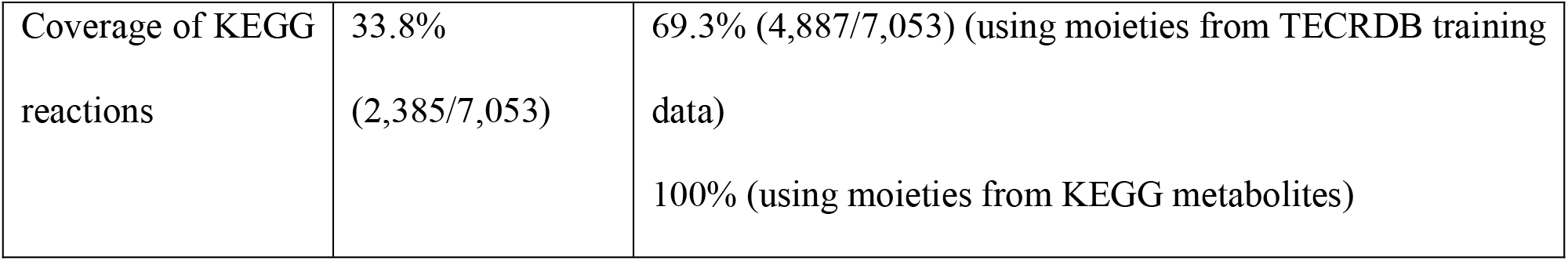
Statistics of 15,278 metabolites and 7,053 reactions from KEGG that can be decomposed into groups/moieties.

|  | GC | dGPredictor |
| --- | --- | --- |
| Coverage of KEGG metabolites | 85.3%<br>(13,032/15,278) | 100% (15,278/15,278) |
| Coverage of KEGG reactions | 33.8%<br>(2,385/7,053) | 69.3% (4,887/7,053) (using moieties from TECRDB training data)<br>100% (using moieties from KEGG metabolites) |

**Fig 3.**
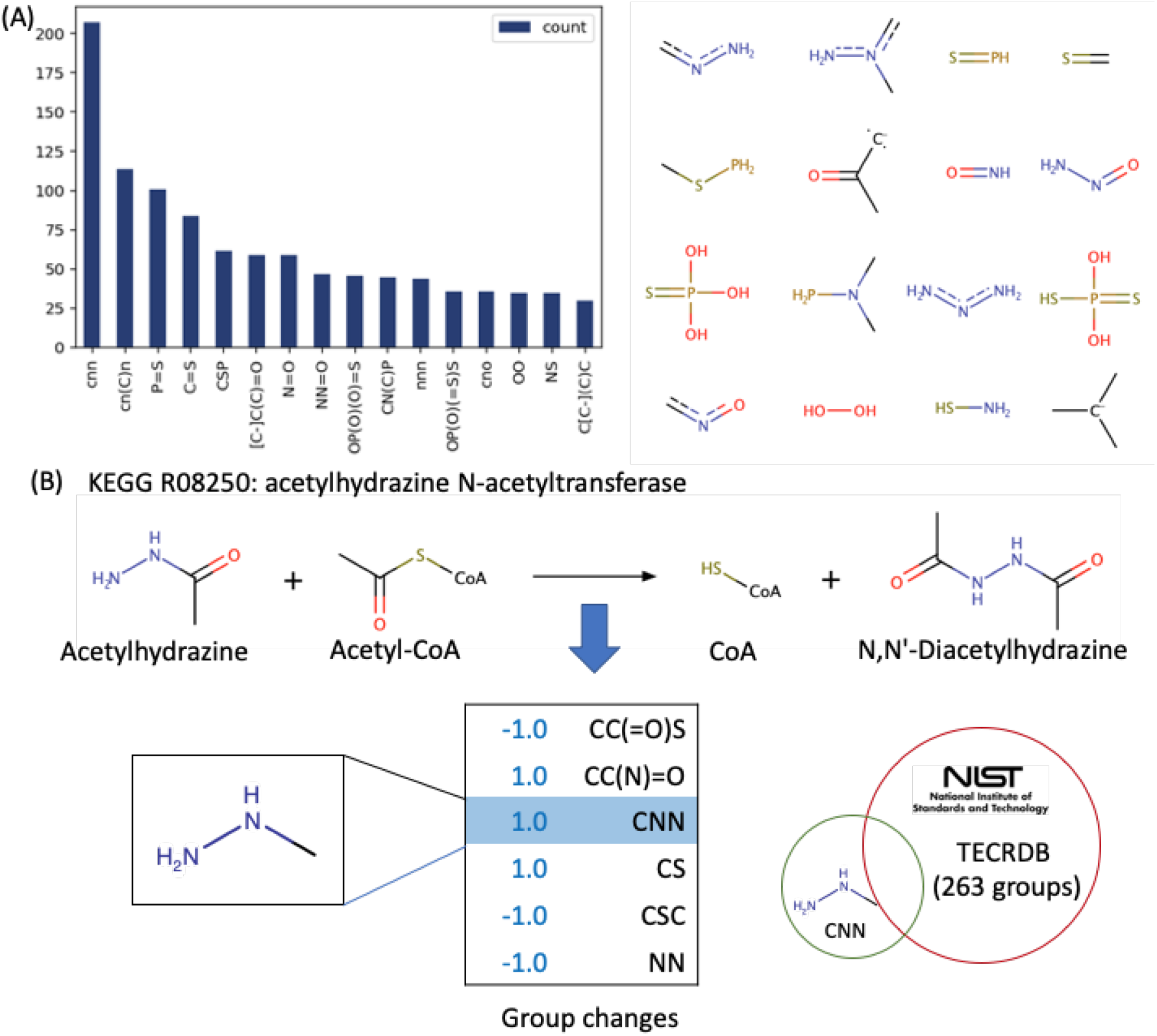
Unique substructure covered by dGPredictor. (A) Unique substructures in the 14.7% metabolites missed by previous GC method. Here the count indicates the occurrence of such substructures in metabolites (B) Example of a reaction with groups (i.e., group “C-N-N”) missing from the 263 groups from TECRDB.

Even though dGPredictor improved the coverage of KEGG metabolites and their corresponding reactions, there are still metabolites with associated moieties absent from the TECRDB dataset, leading to incomplete moiety coverage during model training. For example, the reaction shown in Fig 3B represents six group changes. However, group “C-N-N” is not in the list of the 1,643 moieties generated from metabolites in the TECRDB training dataset (for both radius one and two). To warrant the lack of information in GC, Noor et al. [10] used a standard deviation of *10*^*10*^J/mol to indicate that it cannot estimate Δ_*r*_*G*^*o*^ with any level of certainty. Contrary to the wide confidence interval used in GC, dGPredictor uses Bayesian ridge regression to provide a narrower Δ_*r*_*G*^*o*^ range. Briefly, it assumes an isotropic Gaussian distribution with precision parameter *α* and identity matrix *I* for the standard Gibbs free energy contributions of the groups (Δ_*g*_*G*^*o*^):

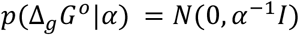

After the hyperparameter optimization during model training (see Methods section ‘Bayesian ridge regression to determine the Gibbs free energy of groups Δ_*g*_*G*^*o*^’), maximizing the log of the posterior distribution in Bayesian ridge regression yields the optimized precision parameters *α* as 9.023 · 10^−4^. The standard deviation of Δ_*g*_*G*^*o*^ in dGPredictor can thus be computed as 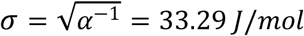. As shown in Fig 2B, Δ_*g*_*G*^*o*^ of 1,643 moieties are well-determined by the normal distribution because the values for 99% of the moieties fall within three standard deviations of the mean (i.e., Δ_*g*_*G*^*o*^ ∈ [−99.87, 99.87]). Therefore, Bayesian ridge regression provides a much narrower confidence interval for all moieties with no prior thermodynamic information. The model predictions can thus increase to 100% reaction coverage with Bayesian regression, although with a wider confidence interval in Δ_*r*_*G*′^*o*^ for reactions that contain moieties without experimental data. Hence, the use of an automated moiety-based description with Bayesian regression allows expanding the set of Gibbs free energy predictions while narrowing the overall confidence interval as compared to the previous GC method.

### dGPredictor enhances prediction scope by accounting for reactions with no GC-defined group changes

A limitation of previous GC methods lies in their inability to predict Δ_*r*_*G*′^*o*^ of reactions with no group changes, which are ultimately assigned a Δ_*r*_*G*′^*o*^ value of zero. These reactions mainly belong to classes such as isomerase and transaminase, which are known to have a Δ_*r*_*G*′^*o*^ far from zero [7]. One such example is the enzymatic reaction catalyzed by GDP-D-mannose 3,5-epimerase (EC number: 5.1.3.18), with an experimentally measured equilibrium constant *K’* = 1.94, implying a Δ_*r*_*G*′^*o*^ of 1.7 kJ/mol [33]. The reactant and product, GDP-mannose and GDP-L-galactose, respectively, are structural isomers (Fig 4B), due to which GC-based methods are unable to capture a group change in the reaction. dGPredictor accounts for the inherent stereochemical changes by capturing the clockwise and anticlockwise configurations of the two chiral centers (Fig 4B). The predicted Δ_*r*_*G*′^*o*^ is 1.74 kJ/mol, which is quite close to the experimental measurement of 1.7 kJ/mol. 319 reactions in the KEGG database are associated with no group changes, as defined by GC-based methods (Fig 4A). Most of them are transferases (EC 2) and isomerases (EC 5), which cannot be captured due to no group changes from the 163 expert-defined groups. dGPredictor described 86.83% of the reactions with no group changes (i.e., 277/319) and, in particular, 39.71% isomerases (i.e., 110/277). This is because the simultaneous inclusion of moieties spanning distances one and two allows us to consider additional details of the localized bonds and atoms, thus alleviating problematic cases of no moiety changes being registered when considering single bonding distance. For example, the moiety description of an aminotransferase reaction (EC 2) (Fig 4C) generates identical moieties for both substrates and products, leading to an empty group change vector when considering bonding distance one moieties. However, allowing an additional bonding distance of two helps dGPredictor generate unique moieties, thus resolving the issue of zero change and in turn leading to a non-zero Δ_*r*_*G*′^*o*^ value.

**Fig 4.**
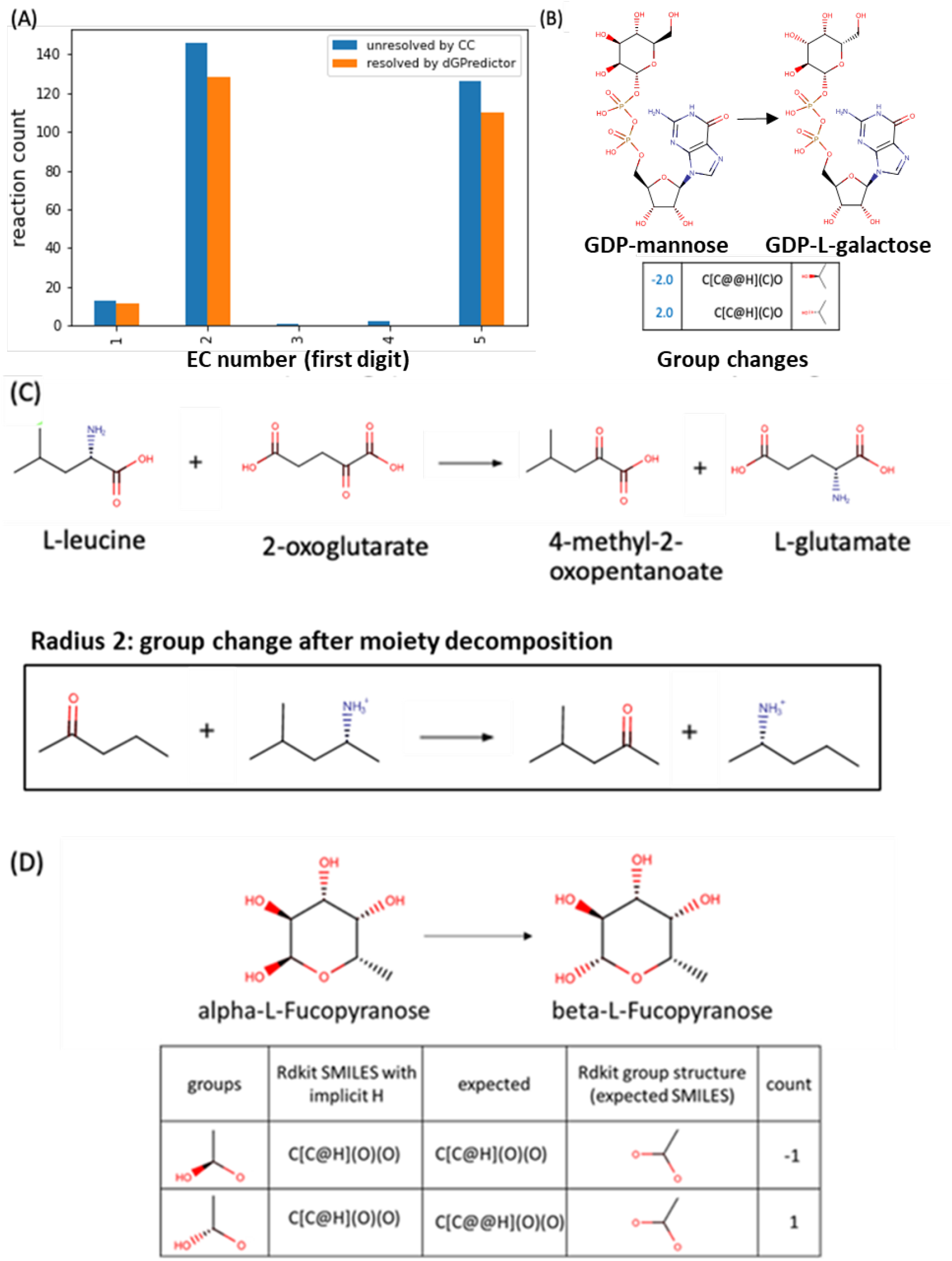
Increased coverage of reactions with no GC-defined group changes. (A) The number of reactions that have no group changes defined by GC (blue) and resolved by dGPredictor (orange), (B) GDP-mannose 3,5-epimerase (R00889) reaction that can be resolved to give group change using stereochemistry information, and examples of reactions in EC 2 (C) and EC 5 (D) that cannot be resolved by dGPredictor using only bonding distance one.

Nevertheless, 42 reactions could still not be resolved because both M1 and M2 descriptions used by dGPredictor were identical between reactants and products. This is because the RDkit tool [34] used for computing moieties stores molecules with implicit hydrogen (i.e., not explicitly present in the molecular graph) and ignores the “C-H” bond when generating the SMILES string. Thus, cannot identify the chiral center when the “C-H” bond is responsible for tetrahedral stereochemistry which leads to same SMILES string for different moieties and a loss of stereochemical information for reactions using M1 moiety change. An example is shown in Fig 4D, where KEGG reaction R10764 that converts *α*-L-Fucopyranose to *β*-L-Fructopyranose is expected to have non-empty group changes because the moieties have different stereo configurations (i.e., [C@H] or [C@@H]) for bonding distance one. In this case, RDkit returning an empty group can be alleviated by combining both distances to capture a non-zero moiety change. This flexible moiety consideration is the primary reason behind dGPredictor providing ∼87% more coverage for reactions with no group/moiety change compared to previous GC methods and the remaining 13% could be tackled using more advanced molecular descriptors such as Neural Graph Fingerprints [15] that account for higher-order interactions instead of local atom/bond information.

### User-friendly interface for Gibbs free energy calculation of novel reactions and metabolites beyond KEGG

Advances in computational pathway design have expanded the range of microbial product synthesis to non-natural synthetic molecules and drug precursors, leveraging broad-substrate range enzymes [35,36] promiscuous activity [37], and even *de novo* enzyme design [38,39]. However, tools such as novoStoic [40] generally treat novel transformations as being reversible, necessitating additional scrutiny to ensure the thermodynamic feasibility of the designed pathway. The ability of dGPredictor to characterize unseen metabolites, and by extension, chemical transformations, accompanied by an automated moiety-based framework enables its use in pathway design tools to eliminate thermodynamically infeasible predictions. In this section, we demonstrate how dGPredictor can predict Δ_*r*_*G*′^*o*^ for novel reactions which do not span known biochemical reactions using chemical moieties, and a GUI that can help users query the same.

The dGPredictor input format uses KEGG IDs to indicate the substrates and products of a reaction. For example, dGPredictor can recognize ‘C00096 <=> C02280’ as the reaction “GDP-mannose <=> GDP-L-galactose” (discussed in above section). However, KEGG is only one of many databases consolidating biochemical reactions, has a lower metabolite content [41], and does not capture novel molecules that often show up in the reactions of retrosynthetic metabolic pathways [42]. Therefore, we designed dGPredictor to allow for user-defined chemical structures as an additional input to estimate Δ_*r*_*G*′^*o*^ for any novel reaction. A user-friendly interface has been developed (https://github.com/maranasgroup/dGPredictor) to facilitate thermodynamic analysis for reactions. Metabolites with known chemical structures can be entered using KEGG IDs and InChI strings are required for molecules not present in the KEGG database. For example, in the *de novo* pathway found by novoStoic for pinosylvin (C01745) degradation, a deoxygenase enzyme catalyzes the first step and produces an aromatic product (see Fig 5). KEGG IDs identify all metabolites except the reduced product ‘N00001’ in the reaction “C01745 + C00004 <=> N00001 + C00003 + C00001”. Here, we use id “N00001” to represent the novel metabolite 3-Phenethyl-phenol, where N refers to novel. The user can indicate that the reaction involves a novel metabolite (i.e., absent in the KEGG database) by clicking a checkbox and filling in the InChI string placeholder under the stoichiometry of the reaction to describe the atom composition and bonds in metabolite N00001 (see Fig 5). Similar to eQuilibrator [11], we allow the customized input of intracellular conditions (i.e., pH and ionic strength) via two sliders. With all the information properly defined, on clicking the “search” button, dGPredictor first show the chemical structures in the reaction and then calculate the standard transformed Gibbs energy *Δ*_*r*_*G*′^*o*^ of the reaction at a particular pH and ionic strength. The output information also displays the standard deviation of predictions from Bayesian ridge regression and the moiety changes as defined by dGPredictor. Note that the standard transformed Gibbs energy *Δ*_*r*_*G*′^*o*^ is not a function of the actual metabolite concentrations. The measured concentrations of metabolites (*c*_*i*_) can be used to calculate the actual Gibbs free energy for a reaction 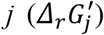 via the equation 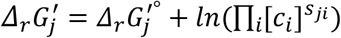 where, *S*_*ji*_ is the stoichiometry of metabolite *i* in reaction *j*. We envision the developed GUI to facilitate easy adoption of dGPredictor to the broader metabolic engineering and synthetic biology community.

**Fig 5.**
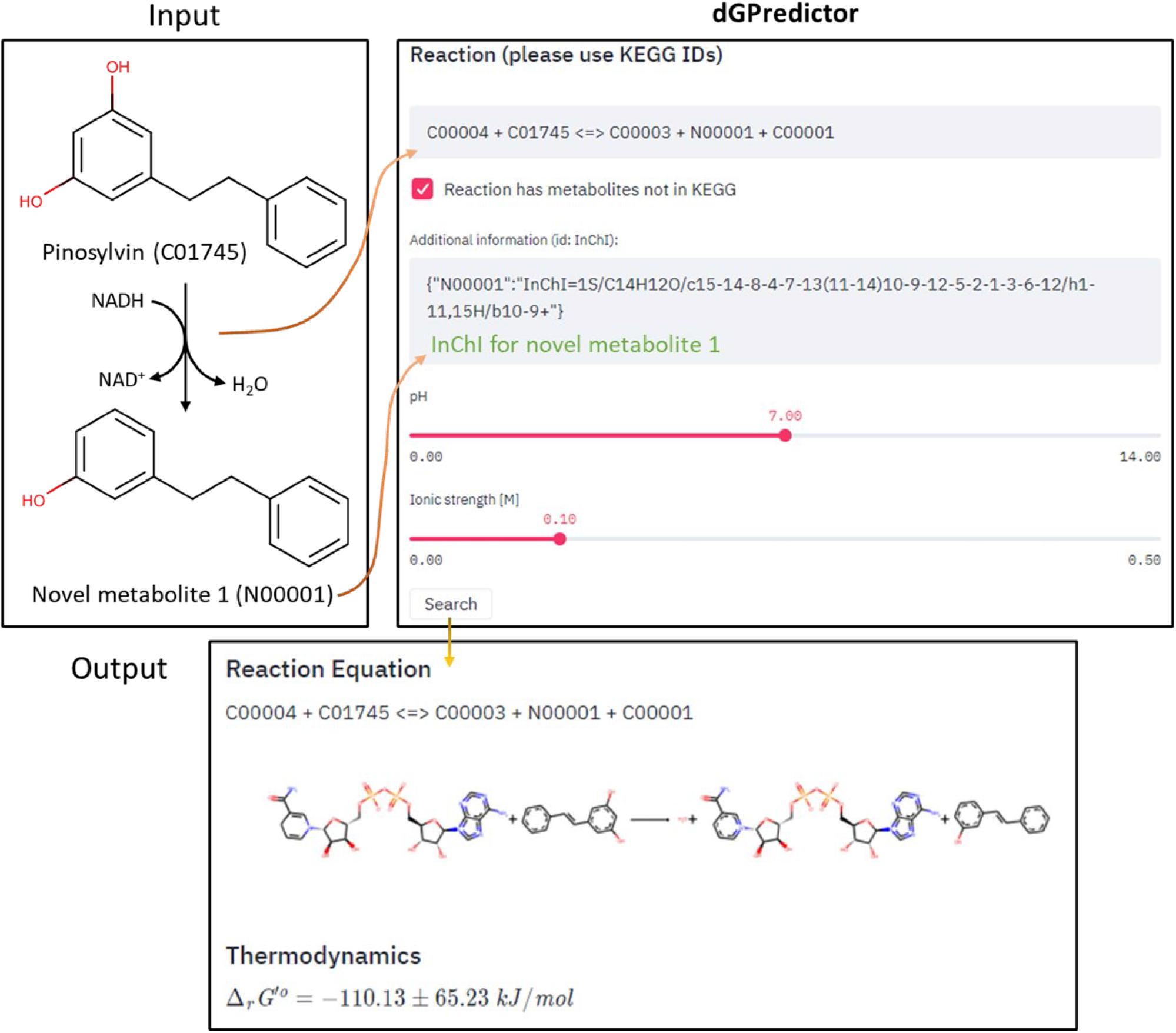
The graphical user interface of dGPredictor. It allows the input of metabolites using their KEGG IDs (for known compounds) and also InChI strings (for novel compounds) in a chemical reaction. Intracellular conditions of pH and ionic strength can also be adjusted using sliders. The final output shows the standard transformed Gibbs energy *Δ*_*r*_*G*′^*o*^ of the reaction at a particular pH and ionic strength.

### Assigning thermodynamics-derived directionality to reaction rules in *de novo* pathway design

As dGPredictor generates automated molecular descriptions, it can be used in conjunction with pathway design tools to ensure that individual reaction, and in turn, the overall pathway is thermodynamically feasible. Pathway design tools such as novoStoic, RetroPath, and RetroPath2.0 [40,43,44] can be integrated with the group change vector of dGPredictor as reaction rules to design *de novo* pathways. We used the novoStoic tool to illustrate this integration and deployed dGPredictor on the 3,603 unique reaction rules generated from KEGG database. We found that based on the predicted standard free energy of change, a number of novel reactions can reliably be flagged as irreversible (i.e., |*Δ*_*r*_*G*′^*o*^| > 20 kJ/mol). This enables the elimination of intermediate steps with a high free energy barrier, thereby significantly reducing the number of candidate pathways to be explored.

Similar to the definition of reaction rules in novoStoic [45], which captures changes in the topology of molecular graphs for a substrate to product conversion [46], dGPredictor assumes that reactions with the same rule undergo same substrate to product change, thereby conforming to identical moiety change. We estimated *Δ*_*r*_*G*′^*o*^ for all 3,603 reaction rules and use ±3 standard deviations to obtain confidence intervals (i.e., 99.7% probability that the free energy estimate is within the calculated interval) (Fig 6A and B). We identified 331 reaction rules with predicted free energy confidence intervals that span only positive values (see Fig 6B) implying that they can only be deployed in the reverse direction. As an additional safeguard, we only applied the irreversibility restriction if the absolute value of the predicted free energy of change |*Δ*_*r*_*G*′^*o*^| exceeds 20 kJ/mol, to account for varying metabolite concentrations that may ultimately tilt reaction directionality [47]. This additional constraint reduces the number of reaction rules that can confidently be treated as irreversible from 331 to 325 (see Fig 6C for EC classification for irreversible reaction rules). Therefore, the dGPredictor framework can help identify the direction *a priori* for irreversible reaction rules and eliminate the thermodynamically infeasible intermediate steps. This allows novoStoic to consider not only the overall thermodynamic feasibility of the pathway but also evaluate every single step.

**Fig 6.**
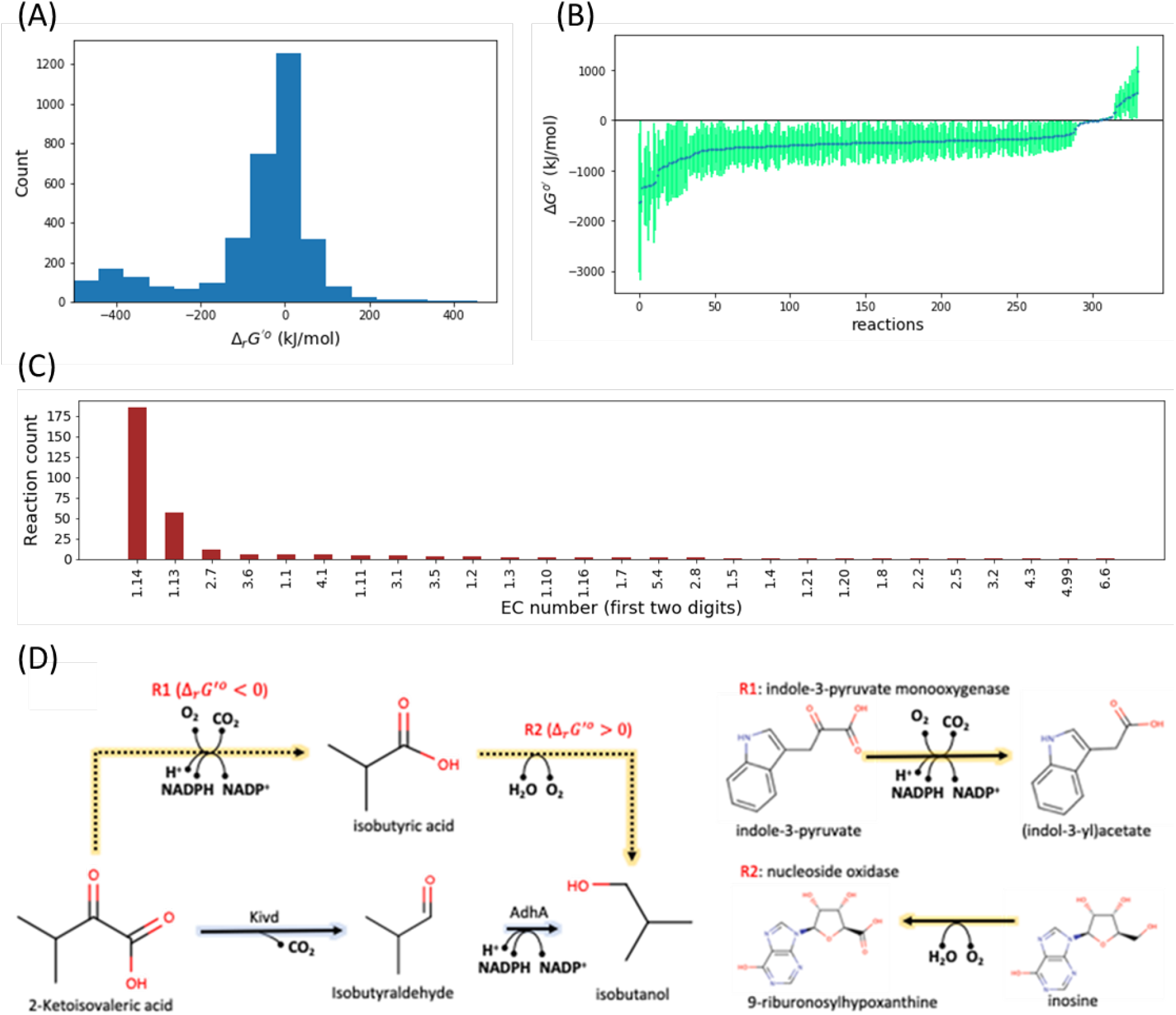
Group change vectors can be used as reaction rules to design *de novo* pathways. (A) *Δ*_*r*_*G*′^*o*^ for all the 3,603 reaction rules, (B) 331 reaction rules found to be irreversible from the uncertainty analysis using ± 3 standard deviations (C) The EC classification for irreversible reaction rules, and (D) an example of thermodynamically infeasible pathways eliminated by using irreversible reaction rules. (see S3 Table for the free energy information of reaction rules and their corresponding KEGG IDs),

To assess the extent of reaction rules’ directionality affecting pathway design predictions, we applied the set of reaction rules to search for isobutanol production pathways (a well-studied product from the Ehrlich pathway [48–51]) using 2-ketoisovalerate as the precursor. Two distinct production pathways were identified by novoStoic (Fig 6D). Pathway A (shown in blue) is the engineered pathway in *E. coli* and *C. thermocellum* with a demonstrated thermodynamic feasibility *in vivo*. Pathway B (shown in orange), however, is thermodynamically infeasible. The first step converts 2-ketoisovalerate to isobutyric acid using a novel reaction R1, which has the same group change as the native reaction of indole-3-pyruvate monooxygenase. However, the second step is a novel reaction R2 similar to the reaction catalyzed by nucleoside oxidase that favors the reverse direction based on dGPredictor thermodynamics analysis. Thus, instead of relying on manual post-processing, direct inclusion of thermodynamic constraints from dGPredictor within the novoStoic pathway design tool can help eliminate infeasible pathways.

## Discussion

This paper presents an automated thermodynamic analysis tool, dGPredictor, based on molecular fingerprints and chemical moieties to estimate the *Δ*_*r*_*G*′^*o*^ of biochemical reactions. Compared with previous group contribution methods, dGPredictor expands thermodynamic calculation coverage to more molecules and reactions with improved accuracy. This is primarily due to the automated fragmentation method that allows incorporating stereochemical information while generating chemical moieties/groups, which was lacking in the group decomposition scheme in previous methods [6,8–10]. The versatility of the fragmentation method is further illustrated by extending predictions to *de novo* reactions, which enables determining the thermodynamical feasibility of synthetic pathways and thus aid strain design algorithms.

First, we showed that the moiety-based automated fragmentation using the SMILES notations can account for stereochemistry in metabolite descriptions. Moieties spanning bonding distances of one and two were generated and used in a linear and non-linear regression framework to ascertain the best performing model. We found that employing a non-linear framework and moieties spanning a bonding distance of two leads to a slight increase in prediction accuracy while being significantly more prone to overfitting. Therefore, an explainable linear model comprising moieties spanning bonding distances of one and two (i.e., M1,2-linear) was determined to be the best performing model and the default for dGPredictor. Notably, the cross-validation MAE (representing overfitting) was comparable to the current state-of-the-art CC method and the MSE (representing prediction accuracy) improved by 78.76% over same training (i.e., the TECRDB database) and cross-validation (i.e., the KEGG database) datasets for a direct comparison of model predictions. We found that the proposed automated fragmentation approach can be implemented on all metabolites in the TECRDB database, unlike the CC method, which uses expert-defined functional groups to decompose metabolites and can only cover 85.3% of metabolites. Molecular descriptions thus obtained were used to estimate Δ_*r*_*G*′^*o*^ for reactions in the KEGG database, leading to an increase in (69.3% vs. 33.8%). However, there remains scope for improvement in the dGPredictor prediction pipeline, which can be aided by increasing the experimental coverage of reactions with experimental Δ_*r*_*G*′^*o*^ estimates, in turn, increasing the number of characterized moieties. dGPredictor can be used to prioritize these experimental targets for Δ_*r*_*G*′^*o*^ estimations by focusing on reactions that comprise unique moieties frequently occur in the uncovered reactions. In addition, quantum chemical calculation [52,53] that *de novo* estimate Δ_*f*_*G*′^*o*^ for metabolites with no experimental measurements and/or novel metabolites generated by retrosynthetic pathway design algorithms can be used to supplement the available experimental data.

A graphical user interface was built for the dGPredictor tool, similar to the group contribution-based web interface eQuilibrator. It can estimate Δ_*r*_*G*′^*o*^ at different intracellular conditions, namely pH and ionic strength. dGPredictor improves upon eQuilibrator by not only estimating the Δ_*r*_*G*′^*o*^ for reactions with compounds in the KEGG database and allowing the user to input novel metabolites using their InChI strings. Thus, our tool broadens the capability of Δ_*r*_*G*′^*o*^ predictions by incorporating novel reactions. This prediction ability is extended to predict the Δ_*r*_*G*′^*o*^ of designed pathways by restricting the directionality to new reaction rules in the *de novo* pathway design tool novoStoic [40]. dGPredictor can help eliminate unnecessary solutions in novoStoic that are not thermodynamically feasible. The moiety change vectors in the dGPredictor can be directly used as allowed reaction rules in novoStoic [45]. dGPredictor thus facilitates an effective search for thermodynamically feasible metabolic pathways by discarding on average 10% of the reaction rules as infeasible, thereby reducing the search space and computational expense for pathway design. dGPredictor can be easily generalized to allow a detailed thermodynamic analysis of large-scale metabolic networks to understand cell metabolism better.

However, it must be noted that there still exists scope for improvement in the molecular descriptors proposed herein. dGPredictor cannot fully resolve reactions with no moiety changes in isomerase reactions due to limitations in the fingerprints defined by RDkit. Therefore, applying advanced 3D molecular descriptors to capture the exact stereochemistry might help capture such reactions. 2D molecular descriptors only contain information for the localized atoms/bonds, whereas a 3D descriptor allows precise capture of molecular shape and interactions [13]. In a recent study, the Quantitative Structure-Activity Relationship (QSAR) method based on Smooth Overlap of atomic position (SOAP) descriptors was shown to successfully capture the 3D atomic environment from conformers [54]. Therefore, utilizing a 3D molecular descriptor for molecule decomposition in the free energy prediction has great potential to account for even more accurate estimation.

## Methods

### Automated fragmentation of metabolites

The algorithm first decomposes the chemical structures of a metabolite into groups of adjacent atoms within a specified bonding distance (i.e., moieties). The bonding distance defines the bond’s proximity to consider from an atom to be described as chemical moieties. We use the InChI string of metabolites as input. Next, we represent each fragment/group with canonical SMILES string. Finally, a group incidence matrix *G*_*i,g*_ is used to represent the count of each moiety as group *g* for every metabolite *i*. We use the automated fragmentation algorithm and represent each group by a unique SMILES string utilizing the Cheminformatic tool RDkit[34] accessed through python.

### Bayesian ridge regression to determine the Gibbs free energy of groups Δ_*g*_*G*^*o*^

Using the molecular descriptions thus obtained, we determine the standard Gibbs free energy contributions of groups Δ_*g*_*G*^*o*^ that allows the best fit of experimental data from TECRDB.

#### Parameters

*SS*: stoichiometric matrix (*R*^*i*×*j*^)

*G*: group incidence matrix that represents group decomposition (*R*^*i*×*g*^)

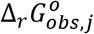 : Gibbs free energy change of reaction j observed in enzyme thermodynamic database TECRDB.

#### Variables

Δ_*g*_*G*^*o*^: standard Gibbs free energy contributions of groups (*R*^*g*×1^)

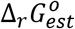 : estimated standard Gibbs free energy change of reactions (*R*^*n*×1^)

The optimization formulation of multi-linear regression with L2 regularization (i.e., ridge regression) to estimate Δ_*g*_*G*^*o*^ by minimizing the sum of the squared estimate of errors (SSE) and the regularization term is shown as follows:

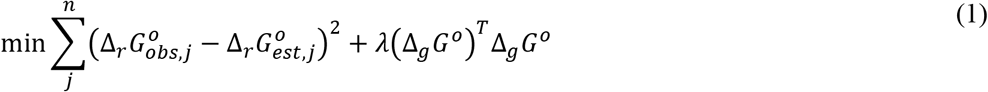

s.t.

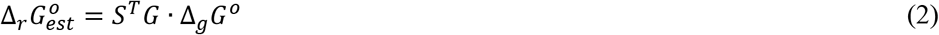

Because the multi-linear regression is prone to overfitting, especially since more groups are defined in our method than previous GC-based methods, L2 regularization 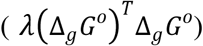 is often applied in ridge regression to reduce cross-validation errors.

Confidence interval analysis from strongly biased models (e.g., ridge regression) relies on bootstrap-based methods. It often produces inaccurate uncertainty estimations [55]. Credible intervals used by Bayesian inference provide an alternative metric to confidence intervals. They have been shown to provide more reliable uncertainty estimation in 13C metabolic flux analysis [56]. Herein, we apply Bayesian ridge regression and define the prior of the Δ_*g*_*G*^*o*^ as the isotropic Gaussian distributions with precision parameter

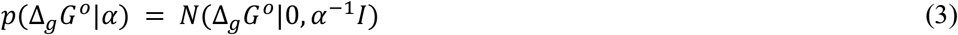

The likelihood function for the 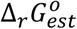 is defined as a Gaussian distribution with a precision parameter *β* :

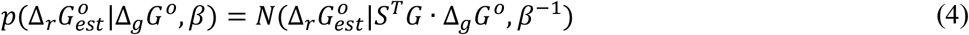

Based on the prior and likelihood function, Δ_*g*_*G*^*o*^ is then estimated by the method of maximum a posteriori estimation (MAP) of the log of the posterior distribution:

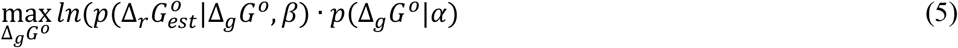

Note that the maximization of the log of the posterior distribution is equivalent to the multi-linear regression with L2 regularization defined in equations (1) and *λ* = *α*/*β* as shown in [25]:

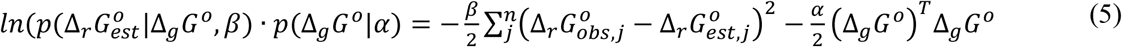

The prediction interval of 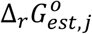 for a new reaction *j* with a group change vector *x*_*j*_ from Bayesian ridge regression can then be calculated as:

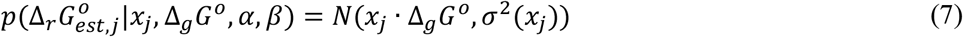

where the variance of the prediction interval can be calculated following the derivation in [34] as:

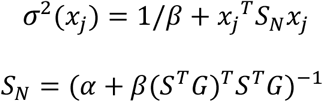

We apply the Bayesian ridge regression function from the scikit-learn python package to train the Bayesian ridge regression model. Scikit-learn optimizes the precision parameters *α* and *β* by iterative re-estimation based on an estimate for “how well-determined” the corresponding Δ_*g*_*G*^*o*^ *i*s by the training data [26]. We found the same parameters and mean squared error in the fitted data upon Bayesian ridge regression to 50 different initial guesses of precision parameters. We infer that the iterative method applied in Bayesian ridge regression in scikit-learn can produce unique, optimized parameters *α* and *β*. Then, the Bayesian ridge regression model can estimate the mean (*x*_*j*_ · Δ_*g*_*G*^*o*^) and standard deviation 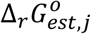.

### Neural networks models for estimating Gibbs free energy of groups Δ_*g*_*G*^*o*^

We choose a feed-forward multi-layer perceptron neural network for nonlinear regression. These networks have neurons that are ordered in layers. The model starts with an input layer (moiety incidence vector), followed by a hidden layer, and ends with an output layer. Fig 7 shows the architecture for a feed-forward neural network. The multi-layer perceptron model maps the moiety incidence matrix of the training data with the output layer using nonlinear functions. The output of the hidden layers is determined by using a rectified linear unit transfer function [57].

**Fig 7.**
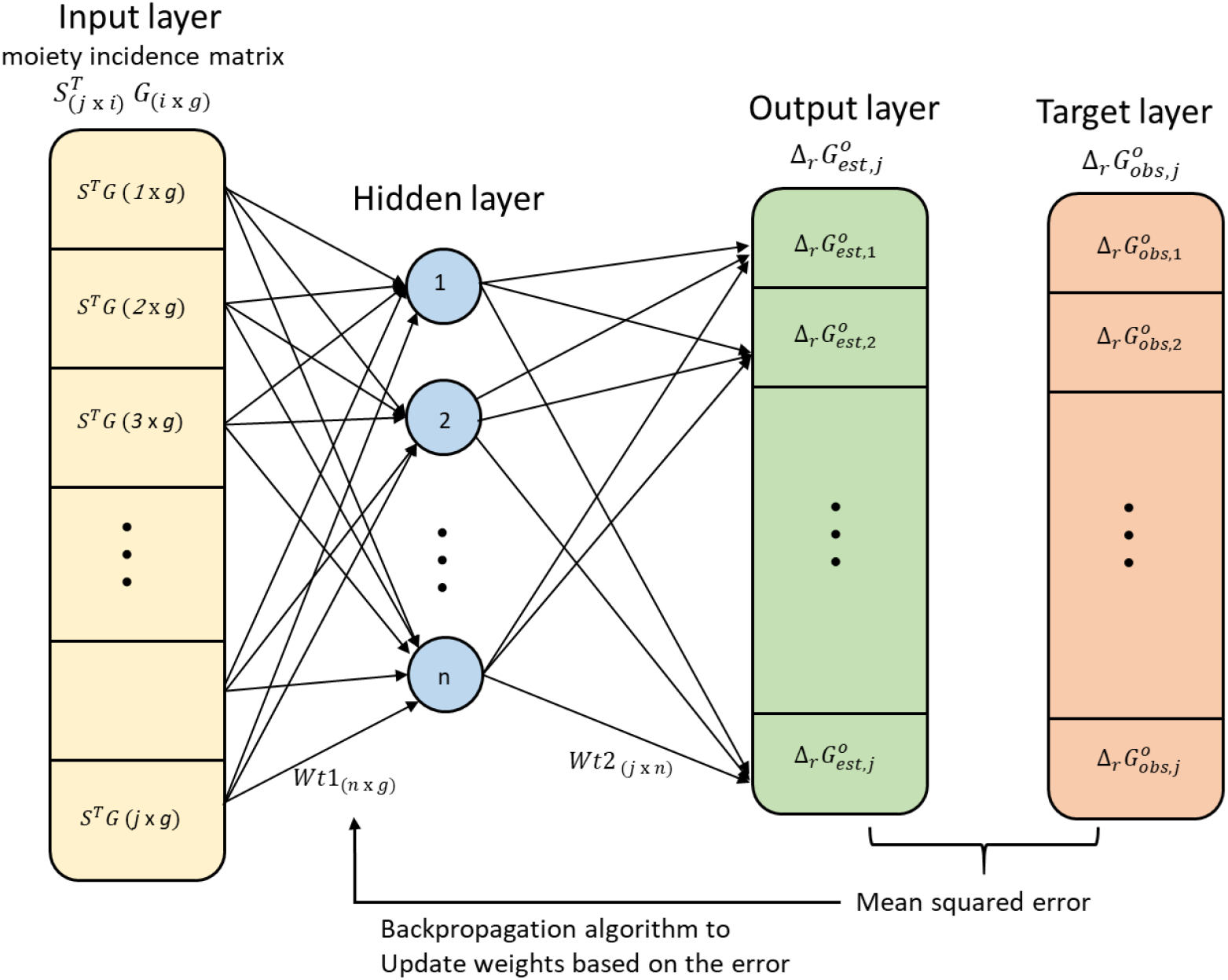
The architecture for multi-layer perceptron neural network. There is an input layer of moiety incidence matrix (*j* × *g* matrix), a hidden layer with n neurons, and an output layer of estimates with j neurons. Here *j* represents the reactions, g represents the groups/moieties, and n represents the neurons in the hidden layer.

We applied the multi-layer perceptron regression from the scikit-learn python package to train the neural network model. LBFGS (Limited memory Broyden-Fletcher-Goldfarb-Shanno) backpropagation algorithm [27] is used to minimize the mean squared error and update the weights of the hidden layers based on the estimates in the output layer. The LBFGS is a faster technique in the family of quasi-newton methods commonly used for parameter estimation in machine learning [27]. Our model has three layers: input, hidden, and output. We considered the scikit-learn package default single hidden layer with 100 neurons to build the neural network model. Studies suggest that a single layer can adequately approximate any function which maps one finite space to another [28]. Scikit-learn package automatically estimates the number of neurons based on the cross-validation results for an accurate fit.

At last, we build three different models for bonding distance one, two, and combining both distances. The M1,2-nonlinear model inputs the moiety incidence matrix for both the radiuses in a single input matrix. The radius one and two have 263 and 1380 unique moieties for 673 metabolites in 4001 reactions. Therefore, the combined-NN model considers a total of 1643 moieties to generate moiety incidence. The same leave-one-out cross-validation was performed as linear regression models for a direct comparison of the model performance.

#### Pseudoisomers

The exact structure of the metabolite inside a cell is typically unknown because it exists as a mixture of pseudoisomers with different protonation states. Pseudoisomers are incorporated into dGPredictor by assuming that the intracellular mixture of pseudoisomers follows the Boltzmann distribution. We use Inverse Legendre Transform [9] to calculate the difference between the standard Gibbs free energy of formation of a compound Δ_*f*_*G*^*o*^ of the major pseduoisomer at pH 7 and the transformed Gibbs free energy of formation of the mixture Δ_*f*_*G*′^*o*^:

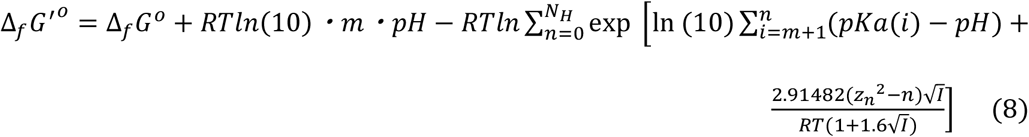

where the acid-base dissociation constant *pKa(i)* of pseudoisomer *i* is calculated using ChemAxon Marvin, *N*_*H*_ is the maximum number protonated hydrogens within a molecule (i.e., the number of *pKas), m* is the number of hydrogens of the reference pseudoisomer, *z*_*n*_ is the total charge the pseudoisomer *n, I* is the ionic strength, *R* is the gas constant, and *T* is the temperature.

The experimental measurement in TECRDB for a reaction for the apparent equilibrium constants *K*′, which is used to calculate the standard transformed Gibbs energy of reaction *j*:

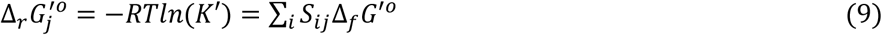

Replacing Δ_*f*_*G*′^*o*^ *in* equation (9) with equation (8), the resulting linear equations can be used to calculate Δ_*f*_*G*^*o*^ using Gaussian elimination. When predicting Δ_*f*_*G*′^*o*^ *of* a reaction, the difference between Δ_*f*_*G*′^*o*^ and Δ_*r*_*G*^*o*^ (i.e., ΔΔ*G* = Δ_*f*_*G*′^*o*^ − Δ_*f*_*G*^*o*^) *of* all the metabolites in the reaction calculated from equation (8) can be added to the Δ_*r*_*G*^*o*^ calculated from regression analysis in above sections to correct the contribution of the pseudoisomer mixture. Both the calculation of Δ_*f*_*G*^*o*^ using Inverse Legendre Transform and ΔΔ*G* are implemented using the functions from the component contribution package (https://github.com/eladnoor/component-contribution).

## Acknowledgments

We thank the input given by Debolina Sarkar for helpful discussions.

## Author Contributions

**Conceptualization:** Lin Wang, Costas D. Maranas.

**Formal analysis:** Lin Wang, Vikas Upadhyay.

**Funding acquisition:** Costas D. Maranas.

**Methodology:** Lin Wang, Vikas Upadhyay.

**Supervision:** Lin Wang, Costas D. Maranas.

**Validation:** Vikas Upadhyay.

**Visualization:** Lin Wang, Vikas Upadhyay.

**Writing-original draft:** Vikas Upadhyay, Lin Wang, Costas D. Maranas.

**Writing-review and editing:** Vikas Upadhyay, Lin Wang, Costas D. Maranas.

## Supporting Information

**S1 Table. The list of moieties generated for bonding distance one for all the TECRDB metabolites**.(TXT)

**S1 Table. The list of moieties generated for bonding distance two for all the TECRDB metabolites**.(TXT)

**S3 Table. The Gibb’s free energy information of 3603 reaction rules and their corresponding KEGG IDs**. (XLSX)

